# Orexin-1 receptor signaling in ventral pallidum regulates demand for the opioid remifentanil

**DOI:** 10.1101/353896

**Authors:** Aida Mohammadkhani, Caroline B. Pantazis, Hannah E. Bowrey, Morgan H. James, Gary Aston-Jones

**Affiliations:** School of Cognitive Sciences (SCS), Institute for Research in Fundamental Sciences (IPM), PO Box 1954851167, Tehran, Iran; Brain Health Institute, Rutgers University, Piscataway, NJ 08854; Save Sight Institute, University of Sydney, NSW Australia 2000; Florey Institute of Neuroscience and Mental Health, University of Melbourne, VIC, Australia 3052

**Author notes:** Corresponding author: Gary Aston-Jones, Ph.D., 683 Hoes Lane West, Piscataway NJ 08854, E, Submitting author: Aida Mohammadkhani.

## Abstract

Signaling at the orexin-1 receptor (Ox1R) is important for motivation for various drugs of abuse. Recently, our laboratory showed that systemic blockade of Ox1Rs decreased motivation for the potent and short-acting opioid remifentanil, as well as remifentanil consumption at null cost (Porter-Stransky et al., 2017). However, the central site(s) through which orexin acts to regulate opioid-seeking behaviors are not known. We investigated ventral pallidum (VP) as a potential site of orexin action, as VP is a known mediator of opioid reward and is densely innervated by orexin-immunoreactive fibers. We used a within-session behavioral economics (BE) paradigm in which remifentanil price (responses/μg iv remifentanil) was sequentially increased throughout the session. Rats were implanted with bilateral cannulae into caudal VP (cVP), through which microinjections of SB334867 (SB), an Ox1R antagonist, were given prior to BE and reinstatement testing. Contrary to systemic SB treatment, intra-cVP SB preferentially reduced motivation (increased demand elasticity and reinstatement) for reminfentanil, without affecting remifentanil consumption at null cost. These effects were specific to cVP, as control microinjections of SB immediately dorsal to cVP did not affect remifentanil-seeking. Baseline demand for remifentanil predicted cued reinstatement of remifentanil seeking and the efficacy with which intra-cVP SB administration reduced motivation for remifentanil. These findings indicate a specific role for cVP Ox1R signaling in mediating the motivational properties of the opioid remifentanil, distinct from its consumption at null cost. Our findings also highlight the BE paradigm as an effective biomarker for predicting opioid addiction behaviors and treatment efficacy.

**Significance Statement:** Abuse of prescription opioids has risen rapidly and continues to be a major health issue. There is significant urgency to understand the neurobiological and behavioral mechanisms underlying opioid addiction. Here we determined the role of caudal ventral pallidum (cVP) orexin signaling in remifentanil seeking using a within-session behavioral economics paradigm. We report that intra-cVP infusions of the orexin-1 receptor antagonist SB334867 (SB) blocked motivated responding for remifentanil as well as cued ‘relapse’ of remifentanil seeking. In contrast, SB intra-cVP did not affect consumption of remifentanil under low-effort conditions. Together, our findings indicate that decreasing orexin-1 receptor signaling in cVP is a potential strategy for the treatment of opioid abuse.

## Introduction

In recent years, abuse of prescription opioids has risen rapidly and has become a major health issue (Compton and Volkow, 2006; Lewis et al., 2017; Volkow and Collins, 2017; Dobbs and Fogger, 2018). Thus, it is urgent to understand the neurobiological and behavioral mechanisms underlying opioid addiction to facilitate development of new treatments for this disorder. The hypothalamic orexin (hypocretin) system is a key modulator of drug seeking behaviors across a variety of drugs of abuse, including opioids (Harris et al., 2005; Mahler et al., 2014a; Baimel et al., 2015; James et al., 2017). Orexin-containing neurons are located in hypothalamus and project widely throughout the brain, targeting two G-protein-coupled receptors (orexin receptors 1 and 2; Ox1R and Ox2R, respectively). In particular, signaling at Ox1R has been shown to mediate motivation and addiction (for review, James et al., 2018). We found that blocking Ox1Rs can disrupt opioid seeking behaviors: systemic blockade of Ox1R signaling reduced breakpoints for heroin in a progressive ratio task (Smith and Aston-Jones, 2012) and attenuated demand (motivation) for remifentanil in a behavioral economics (BE) paradigm (Porter-Stransky et al., 2017). Systemic Ox1R blockade also reduced low-effort (fixed ratio [FR] 1) self-administration of heroin and remifentanil, indicating a potential additional role for the orexin system in mediating the low-effort consumption of opiates. Despite these data, little is known about where in the reward circuitry orexin acts to contribute to opioid reward behavior.

One region of interest is ventral pallidum (VP), as this brain structure is innervated by orexin fibers (Peyron et al., 1998; Baldo et al., 2003) and VP neurons express both Ox1Rs and Ox2Rs (Marcus et al., 2001). Human studies indicate that drug-related cues robustly activate VP (Childress et al., 2008), and inactivation of VP in rats prevents various drug behaviors, including acquisition and expression of learned preferences for environments paired with morphine reward (Dallimore et al., 2006). Moreover, VP inactivation decreases FR5 heroin self-administration (Hubner and Koob, 1990). Interestingly, caudal VP (cVP) contains the greatest density of orexin fibers in the VP (Baldo et al., 2003), and previous studies found that orexin microinjection into this region enhances the hedonic impact of sucrose (Ho and Berridge, 2013). Taken together, these studies indicate a potential role for orexin signaling in cVP in opioid reward.

The present study was designed to test the hypothesis that orexin signaling in cVP regulates demand (motivation or low-cost consumption) for the ultra-short acting opioid remifentanil. We focused on remifentanil because its short half-life lends itself well to BE analysis with the threshold procedure (Glass et al., 1999), and it contributes to prescription opioid abuse (Zacny and Galinkin, 1999; Baylon et al., 2000; Panlilio and Schindler, 2000; Levine and Bryson, 2010). We determined remifentanil demand using a within-session BE paradigm that provides a quantitative measure of both the desired amount of remifentanil at low effort, and the motivation for this drug as the required effort increases. We first sought to confirm the results of our previous study, which found that systemic administration of the selective Ox1R antagonist SB334867 (SB) reduced motivation (increased demand elasticity; α), and reduced free consumption (Q_0_), of remifentanil. We then compared these effects to those following local microinfusion of SB into cVP. In agreement with our previous study, systemic SB reduced motivation (increased demand elasticity; α) and decreased free consumption of remifentanil (Q_0_). In contrast, intra-cVP SB increased demand elasticity for remifentanil without affecting free consumption, indicating a preferential role for orexin signaling in cVP in motivation for remifentanil. Both systemic and local blockade of Ox1R signaling in cVP also reduced cued reinstatement of remifentanil seeking. Together, our findings point to cVP as a critical site of Ox1R signaling that drives motivation for remifentanil reward, and thus highlight a novel target for therapeutics designed to treat opioid addiction.

## Materials and Methods

### Subjects

Male Sprague Dawley rats (n=24; initial weight 275–300 g; Charles River, Raleigh, USA) were single-housed and maintained under a 12h reverse light/dark cycle (lights off at 08:00h) in a temperature and humidity-controlled animal facility at Rutgers University. Food and water were available ad libitum. All experimental procedures were approved by the Rutgers Institutional Animal Care and Use Committee and were conducted according to the Guide for the Care and Use of Laboratory Animals. Rats were handled daily after a 3-day acclimation period at the facility. All experiments were performed in the rats’ active (dark) phase.

### Drugs

Remifentanil (obtained from the National Institute of Drug Abuse (NIDA) Drug Supply Program, Research Triangle Park, USA) was dissolved in 0.9% sterile saline for intravenous (iv) self-administration. SB334867 (SB), a selective antagonist of the Ox1R was supplied by the NIDA Drug Supply Program and dissolved in sterile artificial cerebrospinal fluid (aCSF) at a concentration of 1mM for VP microinjections, as described in our previous study (Mahler et al., 2013). For systemic administration, SB was suspended in 10% 2-hydroxypropyl-b-cyclodextrin in sterile water and 2% DMSO (Sigma-Aldrich, USA), as in our prior studies (Bentzley and Aston-Jones, 2015; Porter-Stransky et al., 2017; James et al., 2018a; James et al., 2018c). SB (30mg/kg) or vehicle was injected at a volume of 4.0ml/kg i.p. 30min prior to testing. We used this dose of SB because we previously reported that lower doses of SB (10mg/kg) did not affect demand elasticity, free consumption, or reinstatement responding for remifentanil (Porter-Stransky et al., 2017). A within-subjects design was used whereby each rat received both SB and vehicle, and the order was counterbalanced across subjects.

### Intravenous catheter and stereotaxic surgery

Rats were implanted with an indwelling catheter into the jugular vein as described previously (McGlinchey et al., 2016). Rats allocated to VP experiments were then placed in a stereotaxic frame (Kopf, Tujunga, USA) and implanted with bilateral stainless steel guide cannulae (22G, 11mm, Plastics One) directed 2mm dorsal to cVP (−0.8mm posterior, ±2.6mm medial–lateral, −7.5mm ventral, relative to bregma (Paxinos and Watson, 1998). Cannulae were secured to the skull using jeweler’s screws and dental acrylic, and stylets were placed into the guide cannula to prevent occlusion.

### Dorsal control and cVP microinfusions

Animals received a mock microinjection the day prior to the first microinfusion test - injection cannulae (28G) were bilaterally inserted into the guide cannula and kept in place for 1 min (no infusions). On test days, rats received microinfusions (0.3μl/side) of either SB (1 mM) or vehicle (aCSF) 5min prior to behavioral testing. Microinjections took place over a 70-sec period, and injectors were left in place for 1min following infusions to limit backflow of the injectate. We have found the dose of SB used here to be effective in other brain areas in prior publications (Harris et al., 2007; Espana et al., 2010; James et al., 2011; Mahler et al., 2013). SB and vehicle microinfusions were counterbalanced in a within-subjects design and administered via polyethylene tubing connected to gastight 10μl Hamilton syringes (Hamilton, USA) set in an infusion pump (Model 975, Harvard Apparatus, USA).

Control microinjections of SB were first carried out 2 mm dorsal to VP to confirm that our observed changes in behavior were not due to dorsal diffusion of SB. This within-subjects control was performed using injectors that projected 0.2mm below the tip of the guide cannula. Dorsal control microinjections were performed in a session prior to VP microinjection sessions.

Each rat received a maximum of 6 VP microinfusions in a counterbalanced within-subjects design. These included microinjections of SB and vehicle for BE performance, cue-induced reinstatement, and locomotor activity.

### Localization of injection sites

Following the final behavioral test, rats were deeply anesthetized with ketamine/xylazine (56.6/8.7mg/kg) and received bilateral microinfusions of pontamine sky blue via the VP infusion cannulae (0.3μl). Brains were flash-frozen in 2-methylbutane, sectioned into 40μm-thick sections, mounted on glass slides, Nissl-stained with neutral red, and cover slipped to localize cannula tract damage and verify injection sites. We sought to test the effect of SB in cVP (AP: -0.1 to -0.8mm relative to bregma), so rats with misplaced VP cannulae that were located in rostral VP (rVP; AP>0.0 mm relative to bregma; n=5) were analyzed as a separate group.

### Self-administration training

The self-administration procedure was similar to a recently published study from our laboratory (Porter-Stransky et al., 2017). All self-administration sessions occurred in standard Med Associates operant chambers. A response on the active lever resulted in remifentanil infusion (1μg over 4s) paired with light and tone cues. After each infusion, a 20s time-out occurred (house light off) when additional presses did not yield remifentanil or cues. Presses on the inactive lever had no consequences. Each session ended after 2h or when 80 infusions were earned, whichever occurred first. Subjects were trained on a fixed ratio-1 (FR1) procedure for a minimum of 6 sessions. After earning a minimum of 25 infusions within 2h for at least 3 consecutive sessions, animals advanced to the behavioral economics procedure.

### Behavioral economics (BE) procedure

Rats were trained to self-administer remifentanil on an FR1 schedule (as described above) before being trained on the BE task, as described previously (Porter-Stransky et al., 2017). Briefly, during the BE task the cost of drug increased by decreasing the amount of drug infused per active lever press every 10min. Therefore, each 110min session tested 11 doses of remifentanil (2, 1, 0.6, 0.3, 0.2, 0.1 0.06, 0.03, 0.02, 0.01 and 0.006 μg/infusion). Each infusion was accompanied by presentation of a light-tone cue; responses on the inactive lever had no consequence. Demand curves were generated for each BE session as previously described (Bentzley et al., 2013; Porter-Stransky et al., 2017) using an exponential demand equation (Hursh and Silberberg, 2008) and least sum of squares approach, to generate estimates of preferred remifentanil intake at zero cost (Q_0_) and demand elasticity (α, the slope of the demand curve). Larger α values indicate greater demand elasticity and are characterized by a greater reduction in responding as drug price increases; this is interpreted as decreased motivation (Bentzley et al., 2013). Smaller α values indicate lower demand elasticity and are symptomatic of continued responding for drug despite increases in cost to obtain drug (increased motivation).

Subjects were trained daily on the BE procedure until stable responding was obtained (≤20% variation in α and Q_0_ over 3 consecutive days) before testing the effects of SB. Between drug treatments, rats were re-tested on the BE procedure for a minimum of 3d and and until behavior returned to pre-treatment levels.

### Extinction and reinstatement tests

After BE testing, rats were subjected to extinction training and locomotor activity testing (described below). The order of extinction training and locomotor tests was counterbalanced. During each 2h extinction session, lever presses yielded neither remifentanil nor cues. Rats received extinction training for at least 7d and until they met the criteria of ≤15 active lever presses for at least two consecutive sessions. Rats were given two 2h cue-induced reinstatement tests with either SB or vehicle (counterbalanced), in which pressing the active lever yielded the cues previously paired with remifentanil infusions, but no remifentanil. Rats received at least 2d of extinction (≤15 active lever presses) between tests. One rat from the systemic SB group died between BE and reinstatement testing, and thus was not included in extinction/reinstatement analyses.

### Locomotor testing

Rats were tested for locomotor activity as described previously (McGlinchey et al., 2016; James et al., 2018b) in locomotor boxes (clear acrylic, 40 x 40 x 30 cm) with Digiscan monitors (AccuScan Instruments). Rats were placed in a chamber for 2h to acclimate to the environment. The next day, rats received a microinjection of either SB or aCSF into cVP 5min before being tested in the locomotor chamber for 2h. Total, horizontal, and vertical locomotor activity were recorded using beam beaks. A within-subjects design was utilized, and drug order was counterbalanced across rats. Rats underwent a one-day washout period in between testing sessions in which they were placed in the locomotor chamber but did not receive drug.

### Data analysis

Analyses were performed in Graphpad Prism V6, except for multiple linear regression analyses which were performed using SPSS Statistics (V19). Acquisition of self-administration, individual differences in BE parameters, and the effects of SB on locomotor activity were analyzed using repeated measures ANOVA and paired t-tests. α and Q_0_ values following SB treatment were transformed to represent percent change relative to their control injections. For VP microinjection experiments, separate analyses were used to compare the effect of SB in cVP and rVP relative to vehicle and dorsal control injections. The effects of SB on active/inactive lever responding during reinstatement tests were assessed using separate 2-way repeated measures ANOVAs with ‘treatment’ (extinction, vehicle, SB) and ‘lever type’ (active, inactive) as the variables. Tukey post-hoc tests were applied to all significant repeated measures ANOVA tests. A linear regression was used to correlate individual Q_0_ and α values. For multiple linear regression analyses, Q_0_ and α values were set as the independent variables, with the dependent variables being active lever responses during cued reinstatement, or the difference in responding between SB and vehicle treatments.

## Results

### Experiment 1: Systemic blockade of Ox1Rs reduced economic demand, free consumption, and cued reinstatement of remifentanil seeking

Figure 1a outlines the timeline of behavioral testing to investigate the effect of systemic treatment with SB on remifentanil seeking behaviors. During FR1 training, rats made more presses on the active lever than the inactive lever (main effect, F(11,131)=3.501, p=0.0003) beginning on the third session (Bonferroni post hoc test, p=0.0173) and throughout the remainder of the training sessions (Fig. 1b). This was associated with an increase in remifentanil infusions across training (one-way repeated measures ANOVA; F(5,65)=11.64, p=0.0003; Fig. 1c). Next, rats were trained on the BE procedure, to determine drug demand at varying costs in a within-session design (Fig. 1d). There was a significant inverse correlation between baseline Q_0_ and α in these BE tests (r =-0.6783, p=0.0311 Fig. 1e), such that individuals with high Q_0_ values exhibited more persistent drug taking as the cost for remifentanil increased than individuals with low Q_0_. We next sought to confirm our previous findings that systemic blockade of Ox1R signaling reduces reminfentanil demand (Porter-Stranksy et al., 2015). This provided a direct comparison to intra-VP experiments in subsequent studies (below). Blockade of Ox1Rs with SB significantly increased demand elasticity (decreased motivation for remifentanil, 30 mg/kg SB versus vehicle, t_9_=2.551, p=0.0312; Fig. 1f). Additionally, this dose of SB reduced remifentanil consumption at low effort (free consumption) compared with vehicle (decreased Q_0_, t_9_=2.382, p=0.0411; Fig. 1g). To test the effect of systemic SB treatment on cued reinstatement behavior, lever pressing was extinguished over a minimum period of 7d such that the number of active lever presses per session was <15 (Fig. 1h). A 2×3 repeated measures ANOVA revealed a significant main effect of treatment (F_2,_ 32=7.720, p=0.0018), and lever (F_2,32_=34.56, p<0.0001), and a significant treatment × lever interaction (F_2,32_=8.672, p=0.001). Post-hoc analyses showed that animals reinstated active lever-pressing to remifentanil-associated cues after vehicle pretreatment (vehicle vs. extinction, p<0.0001), and that this was blocked by systemic administration of SB (SB vs. vehicle, p=0.0005; Fig. 1i). There was no effect of SB (p=0.9441) or vehicle (p=0.9658) pretreatment on inactive lever responding (Fig. 1i).

**Figure 1.**
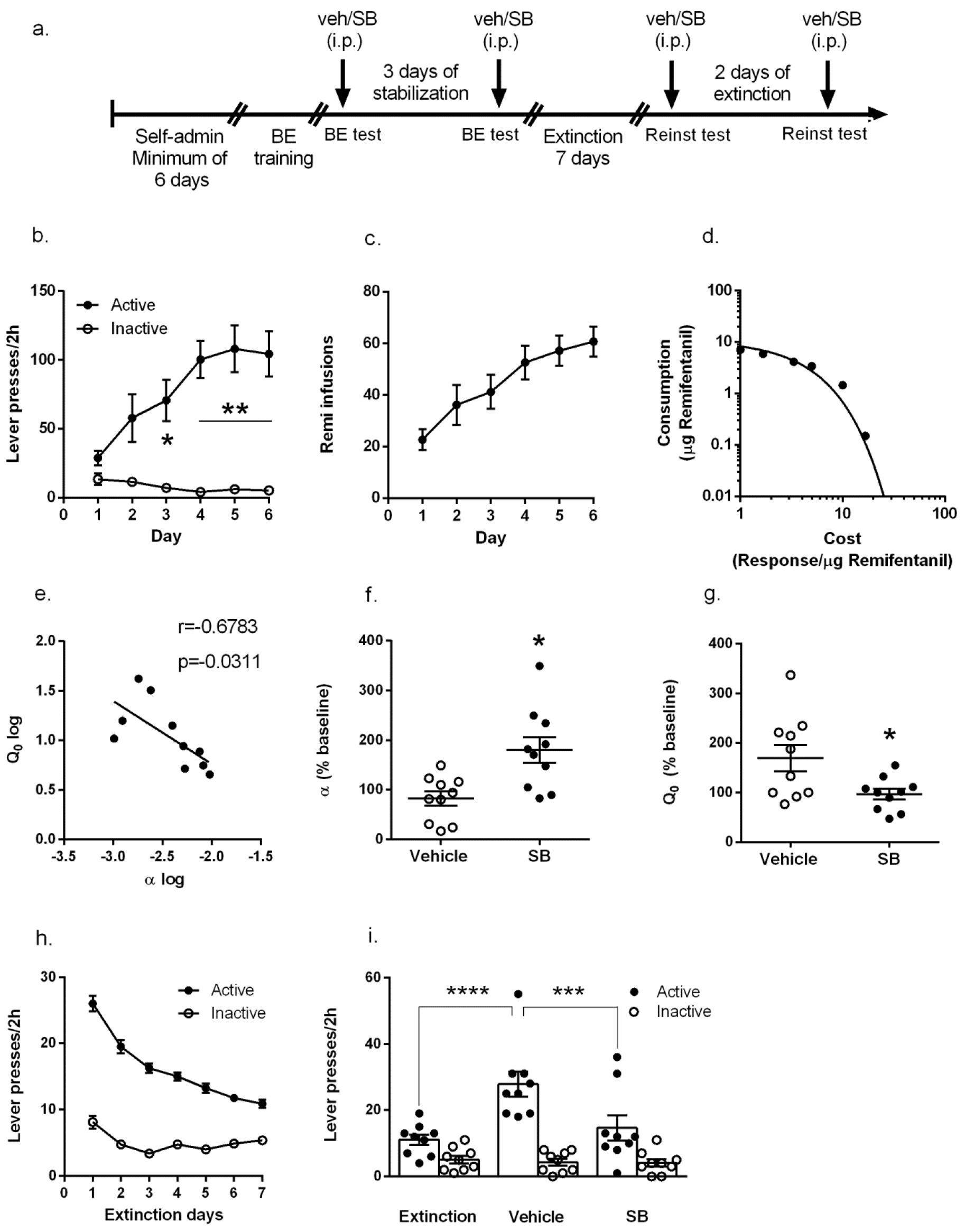
Effects of systemic Ox1R blockade on demand and reinstatement for remifentanil. **(a)** Timeline for behavioral procedure. Rats (n=10) were trained to self-administer remifentanil and then underwent BE training. Lever pressing was extinguished, and animals were then tested in a 2h cue-induced reinstatement session. Arrows in timeline indicate 30 mg/kg SB or vehicle treatments (counterbalanced). **(b-c)** Mean behavioral performance during remifentanil self-administration training. Data reflect the final 6 days of self-administration training. *p<0.05; **p<0.01 active vs. inactive lever. **(d)** A representative demand curve for a single animal during a remifentanil BE session. **(e)** Consumption of remifentanil at null cost (Q_0_) negatively correlated with demand elasticity (α). Each point represents data from a single animal. **(f-g)** SB (30mg/kg, ip) increased α (decreased motivation; f) and reduced Q_0_ (free consumption; g) for remifentanil during BE sessions. **(h)** Mean number of active and inactive lever presses during 7 days of extinction training (n=9). *p<0.05. **(i)** Average active lever presses during the last 3 days of extinction training and 2 h cue-induced reinstatement testing 30 min after pretreatment with 30 mg/kg SB or vehicle. SB had no effect on inactive lever presses. ***p<0.001; ****p<0.0001. Bar graphs represent mean ± standard error of mean (SEM).

### Experiment 2: Antagonism of Ox1Rs in cVP decreased motivation but not free consumption for remifentanil, and decreased cued reinstatement of extinguished remifentanil seeking

A separate group of animals with bilateral cannulae in cVP were used to test cVP as a potential site of action mediating the effects of Ox1R signaling on demand for remifentanil (Fig. 2a). Rats acquired remifentanil self-administration (Fig. 2b,c) and exhibited stable demand (α) for remifentanil that was correlated with Q_0_ values (Fig. 2d) as in Experiment 1. Microinfusion of SB into cVP significantly increased demand elasticity (decreased motivation) compared to microinjections of vehicle (aCSF) or dorsal microinfusions of SB (F_(2, 4)_=15.02, p=0.0012; Fig. 2e). In contrast to systemic injections of SB, intra-cVP microinfusions of SB did not alter remifentanil free consumption (Q_0_; F(2,4)=0.8576, p=0.4358; Fig. 2f). In rats with misplaced SB injections directed at rVP, there was no effect of SB on either demand elasticity (p=0.4878) or free consumption (p=0.8281).

**Figure 2.**
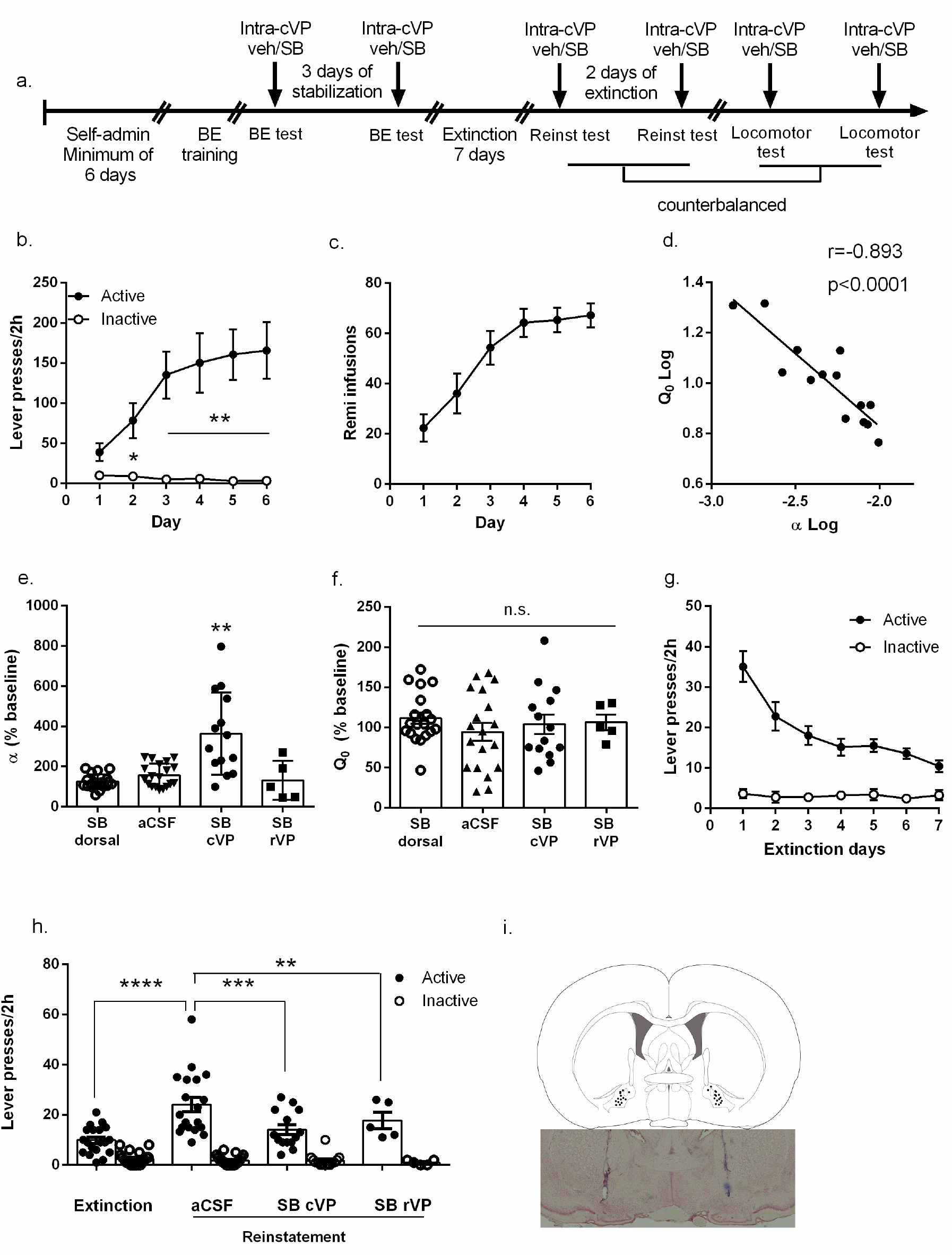
Ox1R antagonism in VP decreases motivation for remifentanil. **(a)** Experimental design of behavioral training and testing, indicating the order and number of injections in VP. **(b-c)** Mean lever pressing during remifentanil FR-1 self-administration (n=14). Data show average number of infusions, and active and inactive lever presses during the final 6 days of FR1 sessions. Rats made significantly more lever presses on the active (drug associated) lever than the inactive lever, beginning on the second session and throughout the remainder of the training sessions. This was associated with an increase in remifentanil (Remi) intake. *p<0.05, **p<0.01 vs. inactive lever. **(d)** There was a significant correlation between α and Qo such that high takers at null effort (high Q_0_) displayed significantly higher motivation (low a) for remifentanil than low takers. **(e)** Demand elasticity (α) was significantly increased (motivation was decreased) following intra-cVP delivery of SB compared to microinfusion of aCSF or SB just dorsal to VP (n=14). No change in α values was observed in animals that received misplaced injections located in rVP (n=5). **p<0.01 compared with dorsal SB. **(f)** Consumption of remifentanil at null cost (Q_0_) was not altered by intra-cVP SB microinfusion, indicating that Ox1R blockade in VP does not alter remifentanil intake at low cost. Similarly, no effect of SB was observed in animals with misplaced (rVP) SB injections (n=5). **(g)** Mean number of active and inactive lever presses during 7 days of extinction training after BE training and testing (n=14). **(h)** Microinfusion of SB into cVP reduced active lever responding during cue-induced reinstatement of remifentanil seeking compared to after aCSF microinjections (n=14). There was also a significant reduction in reinstatement scores in animals with misplaced (rVP) SB injections (n=5). There was no effect of SB vs. aCSF microinjections on inactive lever presses. ***p<0.001; ****p<0.0001. **(i)** Representative schematic (upper panel) and photomicrograph (lower panel; neutral red Nissl stain, frontal section, midline at center) of cannula injection sites from animals that received microinjections in cVP. Injection sites are marked by blue dye microinfused from cannulae at end of experiments. Bilateral injection sites across all rats (black circles; n=14). Bar graphs represent mean ± standard error of mean (SEM). n.s.=not statistically different.

Next we tested the role for Ox1R signaling in cued reinstatement behavior. Lever pressing was extinguished as in Experiment 1 (Fig. 2g). Immediately prior to reinstatement tests, rats were microinjected with either SB or aCSF (counterbalanced) bilaterally into cVP. A 2×3 factor repeated measures ANOVA revealed a significant main effect of treatment (F_(2,_ 52)=8.117, p=0.0009) and lever (F(2,52)=61.38, p<0.0001), and a significant treatment×lever interaction (F(2,52)=8.280, p=0.0008). Tukey’s post-hoc analyses showed that remifentanil-associated cues reinstated active lever pressing after vehicle pretreatment (vehicle vs. extinction, p<0.0001; Fig. 2h). SB in cVP significantly reduced responding on the active lever during reinstatement (SB vs. vehicle, p=0.0008). Separate analyses revealed that SB also reduced reinstatement in animals with misplaced cannulae in rVP (n=5; p=0.0061). There was no effect of SB (p=0.9604) or aCSF (p=0.9955) in cVP on inactive lever responding (Fig. 2h). Together, these data indicate that Ox1R signaling in cVP mediates the motivational properties of remifentanil reward behavior (demand elasticity, cued reinstatement) but is not important for its mediating consumption under null cost conditions.

### Experiment 3: Blocking Ox1Rs in VP did not affect locomotor activity

To test for non-specific effects of intra-cVP microinjections of SB, we assessed general motor activity in an open field apparatus. Bilateral microinfusions of SB into cVP did not alter the total distance traveled (t_13_=0.9848; p=0.3427; Fig. 3a), nor was there an effect on either horizontal (t_13_=1.032; p=0.0754; Fig 3b) or vertical (t_13_=1.436; p=0.1747; Fig. 3c) activity over the 2h test session. These results indicate that effects of SB on demand elasticity and reinstatement were not due to changes in general motoric activity.

**Figure 3.**
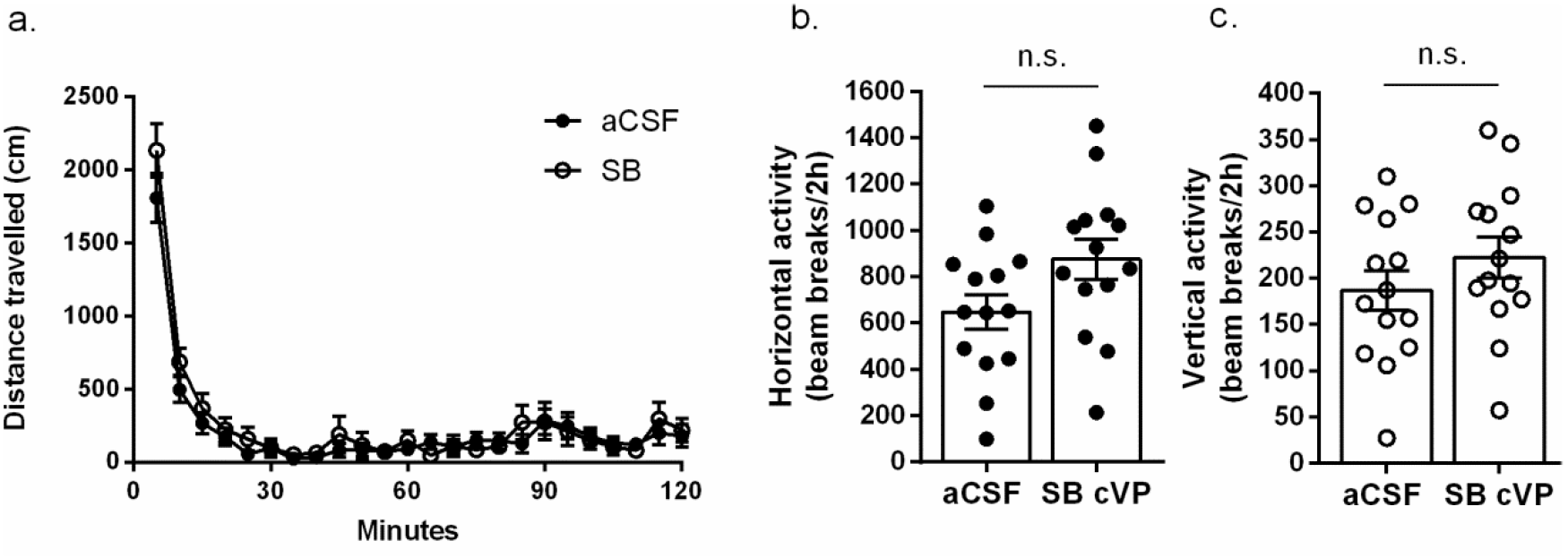
Blockade of Ox1R signaling in cVP had no effect on general locomotor activity. **(a)** Rats displayed no differences in total distance traveled in locomotor chambers after SB microinjection into VP compared to vehicle (n=14). Microinjection of SB into VP had no effect on **(b)** horizontal or **(c)** vertical activity. Bar graphs represent mean ± standard error of mean (SEM). *p< 0.05; ***p<0.001; n.s.=not statistically different.

**Figure 4.**
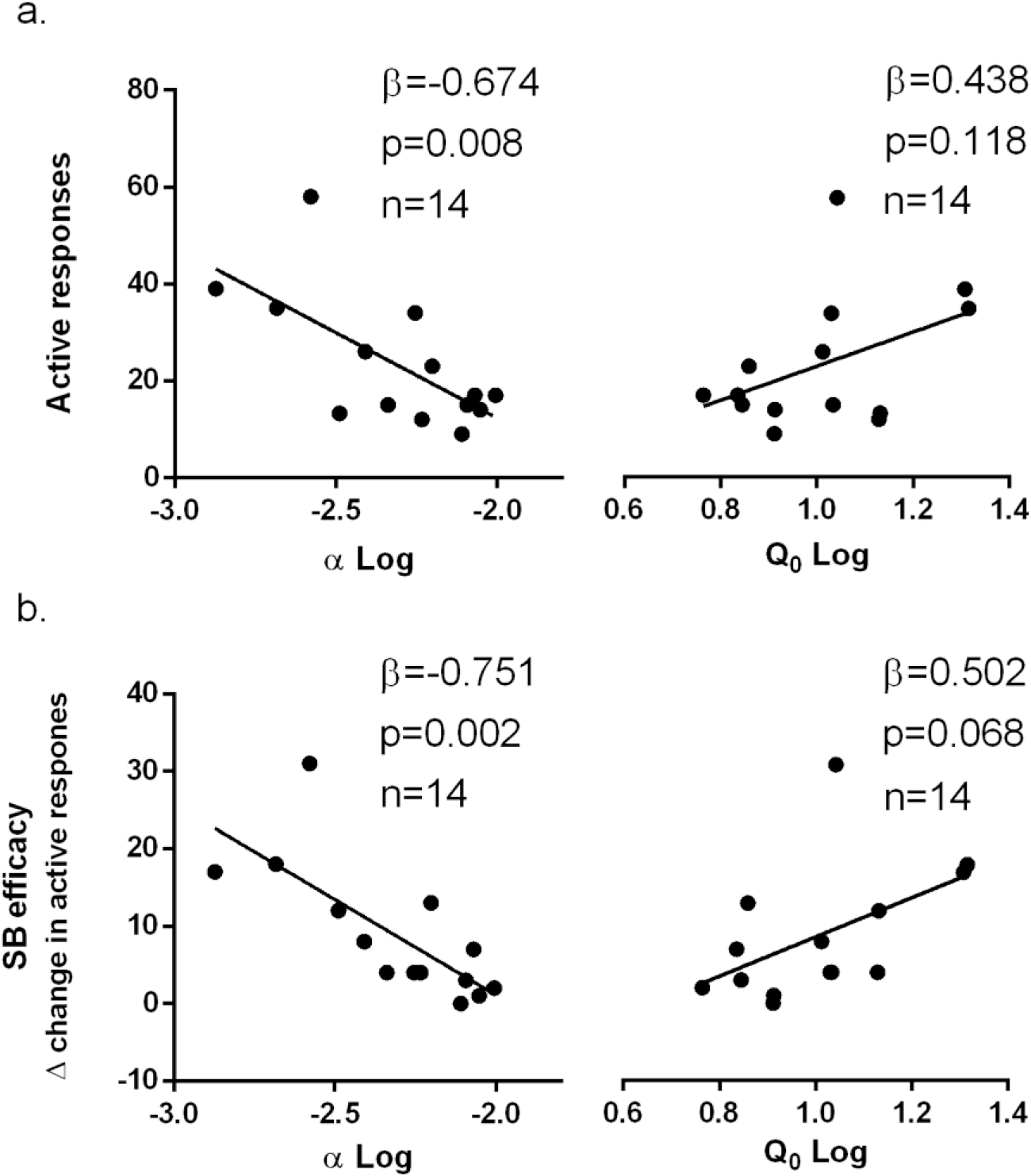
Baseline demand elasticity predicted reinstatement propensity and efficacy of SB microinfusion into VP. (a) Baseline demand elasticity (α), but not free remifentanil consumption (Q_0_), significantly correlated with the magnitude of cue-induced reinstatement with aCSF microinjection, (Q_0_. (b) Baseline demand elasticity (α) correlated with the magnitude of the reduction of cue-induced reinstatement after SB microinjection into cVP. Free remifentanil consumption also correlated with the magnitude of the reduction of cue-induced reinstatement with SB microinjection into VP, but this failed to reach statistical significance (Q_0_). n=14 for all correlations.

### Experiment 4: Baseline demand for reminfentanil predicted cued reinstatement and efficacy of SB

Finally, we sought to test whether ‘baseline’ demand predicted addiction behaviors and SB efficacy, as we have previously reported for cocaine (Bentzley et al., 2014; James et al., 2018a). Multiple linear regression analyses revealed that baseline demand elasticity (α; β=-0.674, p=0.008; Fig. 5a, left), but not free consumption (Q_0;_ β=0.438, p=0.118; Fig. 5a, right), predicted cue-induced reinstatement after aCSF microinfusion into VP. Thus, animals with the lowest α (highest motivation) had greater cued reinstatement behavior. Consistent with previous evidence from our lab indicating that demand elasticity (α) for cocaine can be used to predict those subjects most likely to benefit from the anti-motivational effects of SB treatment (James et al., 2018a), we found here that baseline α (β=-0.751, p=0.002) predicted the extent of reduction in reinstatement by intra-cVP SB (Fig 5b, left). A similar, but non-significant, trend was observed for Q_0_ (β=-0.502, p=0.068; Fig. 5b, right). Together, these data indicate that ‘baseline’ demand elasticity for reminfentanil is a behavioral biomarker for relapse propensity and can be used to predict those individuals most likely to benefit from treatment with an Ox1R antagonist to block reinstatement behavior.

## Discussion

We confirm previous findings (Porter-Stransky et al., 2017) that systemic blockade of Ox1Rs with SB increased demand elasticity (decreased motivation) for remifentanil (α), decreased free remifentanil consumption (Q_0_), and attenuated cue-induced reinstatement of drug seeking. We extend these findings to show that motivation and reinstatement, but not ‘free consumption’, for remifentanil are mediated by Ox1R signaling in cVP. Finally, we show that rats’ baseline remifentanil demand elasticity predicted cue-induced reinstatement responding and the extent of SB effects in cVP on reinstated drug seeking. Together, these results indicate a specific role for orexin signaling in cVP in motivated responding for the opioid remifentanil and highlight demand elasticity as a behavioral biomarker to predict relapse vulnerability and the efficacy of cVP Ox1R antagonism in blocking reminfentanil seeking behavior.

A clear role has been established for orexin signaling in drug seeking behavior (Harris et al., 2005; Mahler et al., 2012; James et al., 2017). Signaling at the Ox1R preferentially mediates high effort drug seeking/taking, or when drug motivation is augmented by external stimuli, such as drug-associated stimuli (Harris et al., 2005; Smith and Aston-Jones, 2012; Porter-Stransky et al., 2017). We previously found that this role extends to opioids, as systemic SB reduced motivated responding for both heroin (Smith and Aston-Jones, 2012) and remifentanil (Porter-Stransky et al., 2017). In addition, Fos activation of lateral hypothalamus (LH) orexin neurons is strongly correlated with the degree of morphine seeking in a conditioned place preference paradigm (Harris et al., 2005). However, unlike with other drugs of abuse, these studies highlighted a role for orexin signaling in free consumption of opioids, as systemic SB treatment reduced FR1 responding for heroin and low-cost consumption (Q_0_) for remifentanil. Here, we confirm a role for Ox1R signaling in both motivation and free consumption of remifentanil reward by showing that systemic SB treatment both increased α and decreased Q_0_ values in the same BE test session.

We investigated cVP as a potential central site of action for SB in mediating remifentanil seeking behaviors, as this region has been implicated in both motivational and hedonic processing. Intra-cVP microinfusions of SB increased demand elasticity (α) but had no effect on remifentanil intake at null cost (Q_0_). These effects were specific to SB in cVP, as we observed no effect of SB microinfusions dorsal to VP and in animals with misplaced injections. VP is a key component of the brain circuitry regulating effort-related choice behavior, as VP inactivation reduces willingness to work on an instrumental task to obtain sucrose reward (Farrar et al., 2008). cVP receives moderate orexin input and expresses a high density of Ox1Rs (Marcus et al., 2001; Baldo et al., 2003), indicating that this region may be a site for orexin modulation of reward behavior. Interestingly however, there is evidence that orexin signaling in VP also mediates the hedonic properties of reward. Microinjections of orexin-A into cVP enhanced the hedonic responses (‘liking’) for sucrose, as assessed by an affective taste reactivity test (Ho and Berridge, 2013). While we have previously argued that Q_0_ might reflect an animal’s hedonic set point for drug (Bentzley et al., 2014), this is difficult to test directly. Thus, we are unable to directly draw conclusions from our data regarding the role for cVP Ox1R signaling and hedonic processing for remifentanil reward. It is unlikely that the lack of effect of intra-VP SB on Q_0_ is due to the dose of SB used, as this dose produces robust behavioral effects in other paradigms (Harris et al., 2007; Mahler et al., 2013) and higher doses of SB would likely have off-target effects (Porter et al., 2001; Smart et al., 2001). It is possible, though, that the previously reported effects of orexin-A infusions in cVP on ‘liking’ for sucrose may be mediated by actions at Ox2R, as orexin-A has similar affinity for Ox1R and Ox2R (Sakurai et al., 1998), and Ox2Rs are expressed in cVP (Marcus et al., 2001). It is also possible that the orexin system plays distinct roles in mediating the hedonic aspects of sucrose versus opioid reward. Clearly, further studies are required to test these possibilities.

Systemic SB is highly effective at reducing cued reinstatement of extinguished drug seeking (Smith et al., 2009; Smith and Aston-Jones, 2012; Plaza-Zabala et al., 2013; Moorman et al., 2017). Such findings led us to hypothesize that the orexin system provides motivational activation by reward-associated stimuli (Mahler et al., 2014a; James et al., 2017). Consistent with this view, we found that both systemic and intra-cVP SB reduced cued reinstatement of extinguished remifentanil seeking. Importantly, VP is part of a broader reward network that underlies reinstatement behavior (McFarland and Kalivas, 2001; Prasad and McNally, 2016); thus, orexin may act at other sites to contribute to cued reinstatement of remifentanil seeking (James and Dayas, 2013; Qi et al., 2013; Matzeu et al., 2018). It will therefore be important that future studies seek to identify the relative contribution of orexin signaling in VP versus other key reward regions in reinstatement of opioid seeking.

Importantly, here we focused on cVP, as this area is more densely innervated by orexin terminals than rVP. However, rVP has also been implicated in motivated reward seeking (Root et al., 2015), including for drugs of abuse (Mahler and Aston-Jones, 2012; Mahler et al., 2014b). Interestingly, in a small number of animals (n=5) with misplaced injections directed at rVP, we observed no effect of SB on BE parameters, but a significant effect of SB on cued reinstatement. This may indicate a functional distinction between cVP vs rVP, where motivation is more associated with orexin signaling in cVP, whereas stimulus-driven seeking might involve orexin input to both regions. However, given the relatively small number of animals in this group, it is difficult to draw comprehensive conclusions from these observations, and it will be necessary that future studies examine this further.

It is possible that blocking Ox1R signaling in our experiments impacted general motor behavior, owing to the role for orexin signaling in arousal (Tsujino and Sakurai, 2013; Li et al., 2014; Li et al., 2017). However we previously reported systemic SB only modestly reduced general locomotor activity in rats with a history of remifentanil self-administration, and that the magnitude of this reduction was not related to SB’s effects on α or Q_0_ (Porter-Stransky et al., 2017). VP has been shown to be critical for the modulation of locomotor activation (Swerdlow et al., 1984; Williams and Herberg, 1987). However, in locomotor activity tests we observed no differences following microinjections of SB in cVP. Moreover, intra-cVP SB did not attenuate inactive lever presses on BE or cued reinstatement tests, and other studies have failed to observe non-specific motor deficits following local injections of SB into other reward regions (Borgland et al., 2009; James et al., 2011; Mahler et al., 2013). We also found that intra-cVP SB had no effect on low effort responding (Q_0_) for remifentanil, further supporting our conclusion that intra-VP SB-induced decreases in demand were not due to motor deficits.

The orexin system is important for motivated responding for natural rewards (Cason and Aston-Jones, 2013b, a) and thus it is possible that the effect of intra-cVP infusions on motivation for remifentanil would also extend to motivated food seeking. Importantly however, evidence indicates that doses of SB that are sufficient to block drug seeking (such as those used here) have no effect on regular chow intake in ad-libitum fed animals (Borgland et al., 2009; Choi et al., 2010; Alcaraz-Iborra et al., 2014), indicating that any potential orexin-based therapy designed to reduce craving and motivation for opioid reward may not interfere with homeostatic feeding.

We previously reported that baseline demand elasticity is a strong predictor of several addiction-relevant endophenotypes (Bentzley et al., 2014; James et al., 2018a). Moreover, we showed that SB is most effective at reducing motivation for cocaine and alcohol in highly motivated animals (Lopez et al., 2016; Moorman et al., 2017; James et al., 2018a; James et al., 2018c). Here, we show that animals with high baseline motivation for remifentanil also have strong cued reinstatement behavior. We also show that high motivation animals exhibited the largest reduction in reinstatement after SB injection, indicating that demand elasticity might also be useful for identifying those individuals most likely to benefit from the therapeutic effects of Ox1R blockade. Given that the BE paradigm can be readily applied in clinical populations (MacKillop et al., 2018), our findings point to the potential utility of this approach for identifying those individuals most likely to relapse following abstinence, and to benefit from pharmacotherapies that interfere with orexin signaling.

In conclusion, our results add to a growing body of literature implicating the orexin system as a target for treating opioid abuse and provide new insights into the corresponding neural substrates. We also show that BE provides a predictive behavioral marker of remifentanil seeking behaviors and the anti-drug seeking efficacy of SB, and thus could be applied clinically to determine opioid addiction liability and optimize pharmacological treatment strategies.

## Acknowledgements

We would like to thank Drs. Kirsten Porter-Stranksy and Brandon Bentzley for their assistance with the remifentanil demand protocol. We would also like to thank Dr. Jennifer Fragale for her helpful guidance in interpreting the data, as well as Nupur Jain for her assistance.

**Funding and Disclosure:**
This work was supported by financial support from the Institute for Research in Fundamental Sciences (IPM; AM), C.J. Martin Fellowships from the National Health and Medical Research Council of Australia to MHJ (No. 1072706) and HEB (No. 1128089), a U.S. Public Health Service award from the National Institute of Drug Abuse to GAJ (R01 DA006214), a NIH F31 grant to CBP (DA042588) and by the Charlotte and Murray Strongwater Endowment for Neuroscience and Brain Health (GAJ).

